# Modeling the dynamics of actin and myosin during bleb stabilization

**DOI:** 10.1101/2023.10.26.564082

**Authors:** Emmanuel Asante-Asamani, Mackenzie Dalton, Derrick Brazill, Wanda Strychalski

**Affiliations:** Department of Mathematics, Clarkson University, Clarkson, Potsdam, NY 13699; Department of Biology, Hunter College, New York, NY 10065; Department of Mathematics, Applied Mathematics, and Statistics, Case Western Reserve University, Cleveland, OH 44106

**Keywords:** Blebbing, *Dictyostelium discoideum*, Cell Migration, Myosin II, Cell cortex

## Abstract

The actin cortex is very dynamic during migration of eukaryotes. In cells that use blebs as leading-edge protrusions, the cortex reforms beneath the cell membrane (bleb cortex) and completely disassembles at the site of bleb initiation. Remnants of the actin cortex at the site of bleb nucleation are referred to as the actin scar. We refer to the combined process of cortex reformation along with the degradation of the actin scar during bleb-based cell migration as *bleb stabilization*. The molecular factors that regulate the dynamic reorganization of the cortex are not fully understood. Myosin motor protein activity has been shown to be necessary for blebbing, with its major role associated with pressure generation to drive bleb expansion. Here, we examine the role of myosin in regulating cortex dynamics during bleb stabilization. Analysis of microscopy data from protein localization experiments in *Dictyostelium discoideum* cells reveals a rapid formation of the bleb’s cortex with a delay in myosin accumulation. In the degrading actin scar, myosin is observed to accumulate before active degradation of the cortex begins. Through a combination of mathematical modeling and data fitting, we identify that myosin helps regulate the equilibrium concentration of actin in the bleb cortex during its reformation by increasing its dissasembly rate. Our modeling and analysis also suggests that cortex degradation is driven primarily by an exponential decrease in actin assembly rate rather than increased myosin activity. We attribute the decrease in actin assembly to the separation of the cell membrane from the cortex after bleb nucleation.

## 1 Introduction

The cortex is the thin layer of the actin cytoskeleton near the cell membrane that gives an animal cell its shape and structure. The highly dynamic nature of actin and actin-associated proteins that constitute the cortex allow the cell to accomplish critical functions, such as cell motility, shaping the extracellular matrix, and cytokinesis [1, 2, 3, 4]. During motility-driven cellular blebbing, the cell membrane detaches from the actin cortex, expanding into a spherical cap (Fig. 1a). A new cortex (bleb cortex) rapidly reforms beneath the membrane (Fig. 1b). During this process, the cortex near the bleb initiation site (Fig. 1a) degrades (Fig. 1c) [5, 6]. We refer to this portion of the cortex as the actin scar. Blebs can be used as a leading-edge protrusion during cell migration, particularly in confined and 3D environments [7, 8]. Blebs also provide an ideal system for studying the assembly, structure, and composition of the cortical actin network [2, 9, 6, 10].

**Figure 1:**
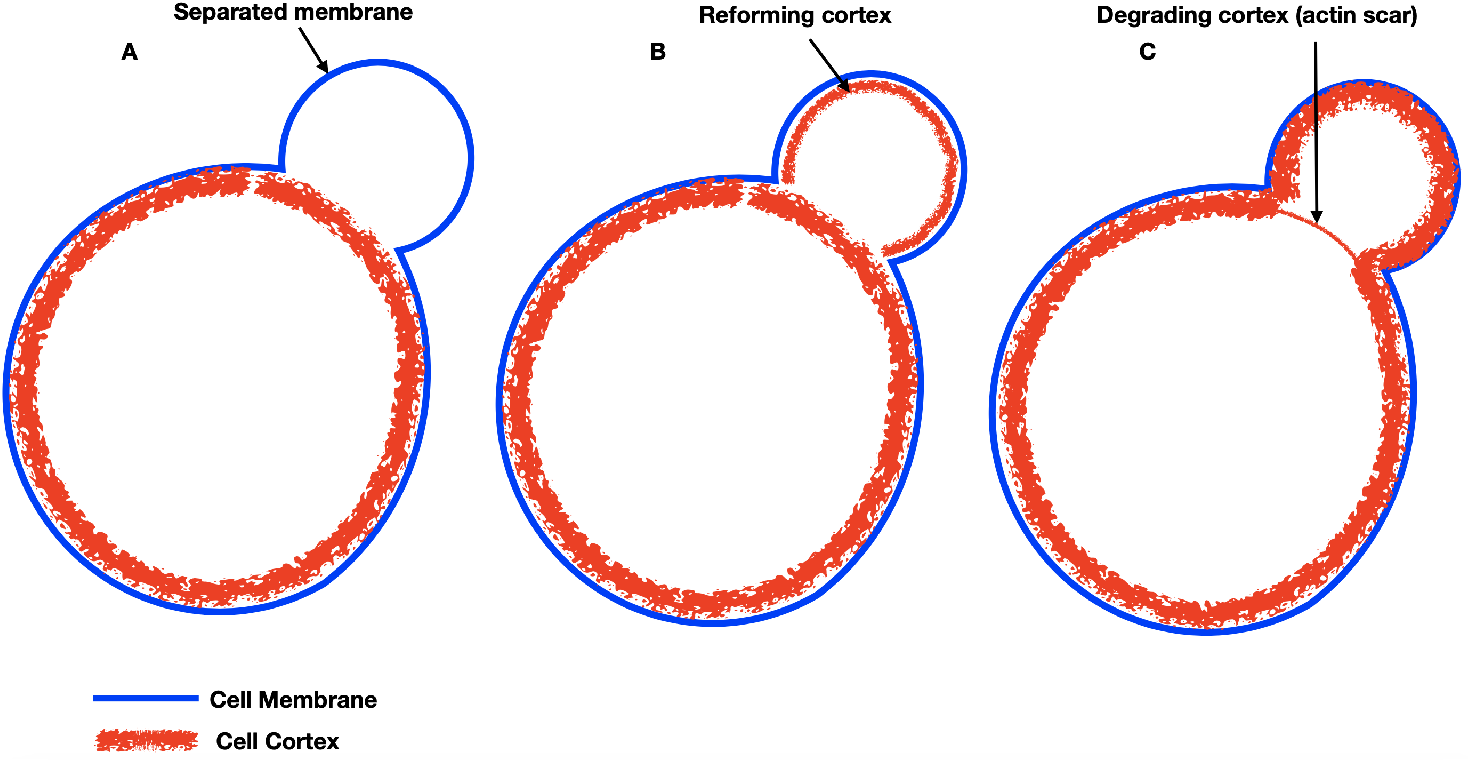
Bleb formation cycle. A) Cell membrane separates from the cortex and expands into a spherical cap. B) A new cortex reforms beneath the protruded membrane C) The old cortex (actin scar) is actively degraded

Blebs can be initiated by cortical rupture, such a by laser ablation in [11], or by a local disassociation of membrane-cortex linker proteins [5]. Recent experimental images of *Dictyostelium discoideum* cells revealed gaps in the cortical actin network near the bleb prior to bleb expansion, pointing to cortex rupture as an intrinsic cell-driven bleb nucleation mechanism [12]. It has been suggested that the myosin-driven contractility contributes to rupture of the cortex [13, 14, 15]. This is supported by recent experiments showing clusters of myosin in the blebbing region of *D. discoideum* [12]. Myosin has also been implicated in generating the excess hydrostatic pressure necessary to expand the bleb [16, 17, 18, 19, 5, 20]. Beyond facilitating bleb nucleation and expansion, little is known about the dynamics of actin and myosin during bleb stabilization in motile blebbing cells. In particular, the role of actin-bound myosin in stabilizing a bleb during cell migration is unknown.

Several studies have reported on actin and myosin dynamics in nonmotile blebbing cells in order to understand assembly of the actin cortex [5, 2]. In [9], the authors measured the timing of protein recruitment in a bleb. First ezrin, an actin cross-linking protein was recruited, followed by actin, actin-bundling proteins, and lastly contractile proteins such as myosin regulatory light chain and myosin heavy chain. Accumulation of actin and myosin during the course of bleb expansion and retraction was reported in [2]. The accumulation speed appeared linear until bleb retraction, when actin and myosin achieved a local peak before decreasing over a period of 100 s. Myosin intensity over time appeared to increase linearly during bleb retraction over a period of 50 s in filamin-deficient macrophages [6]. In motile cells, our group recently reported an increase in myosin intensity during cortex degradation in the actin scar of migrating *D. discoideum* cells, suggesting that myosin assists in this process [12]. However, a thorough quantitative study of the dynamics of the two proteins to support this hypothesis was not undertaken.

In this work, we analyze the temporal dynamics of actin and myosin within the bleb cortex (Fig. 1b) and actin scar (Fig. 1c) of migrating *D. discoideum* cells confined between a glass coverslip and agarose gel. In contrast to previous studies on bleb retraction [5, 6, 11], we consider non-retracting blebs that serve as leading-edge protrusions of migrating cells. Note that if a bleb expands and retracts, the cell does not migrate. We develop mathematical models to support alternative hypotheses about the interaction between actin and myosin and use our data (via parameter estimation) to identify the most likely interaction between the two proteins. Our analysis is based on experimental data, some of which was previously presented in [12]. Our results show that myosin influences the equilibration of the bleb cortex by regulating its disassembly rate but plays a minimal role in cortex degradation. Analysis of our mathematical model reveals that the degradation of the actin scar is driven by an exponential decline in the assembly rate which we postulate is triggered by the separation of the membrane from the cortex. The accumulation of myosin in the actin scar observed in our experimental data is shown to be driven primarily by regulation of its unbinding rates to the cortex, which we suggest is triggered by membrane separation.

## 2 Materials and Methods

### 2.1 Strain and culture conditions

All *D. discoideum* cells were grown axenically in shaking culture in HL5 nutrient medium with glucose (ForMedium) supplemented with 100 *μ*g/mL penicillin and streptomycin (Amresco) at 150 rpm at 22°C. Ax2 cells expressing both LifeAct-RFP and Myosin-GFP were grown in 50 *μ*g/mL Hygromycin B (Thermo fisher Scientific) and 4 *μ*g/mL G418 (Geneticin). The LifeAct-RFP plasmid was obtained from Dr. Chang-Hoon Choi, University of California. The Myosin-GFP plasmids were obtained from Kuwayama Laboratory, University of Tsukuba. Ax2 cell lines were first transformed with Myosin-GFP plasmid. Successful cell lines were then subsequently transformed with LifeAct-RFP. The transformation protocol in [21] was used. In preparation for the under-agarose assay, cells were starved for cAMP competency on 27 mm nitrocellular lose filter pads (GE HealthCare) using the protocol described in [21].

### 2.2 cyclic-AMP under agarose assay

A cAMP under agarose assay was used to induce blebs while visualizing myosin and F-actin localization over a period of 30 seconds. A preheated (90°C, 1 min) number 1 German borosilicate sterile 2 well chamber coverglass slide (ThermoFisher) was loaded with 750 *μ*L of melted 0.7% Omnipur agarose (EMD Millipore). Once solidified, the two wells were created for cells and cAMP and a cAMP gradient established across the wells according to the protocol described in [21].

### 2.3 Image acquisition

Images for examining myosin localization were collected using a Perkin Elmer UltraView ERS spinning disk confocal microscope 103 (Perkin Elmer) equipped with 491 nm and 561 nm lasers (Spectra Services Inc.), as above, which were used to excite myosin-GFP and LifeAct-RFP, respectively. A bright field light source was used to illuminate the membrane. Data was collected at a frame rate of 2.13 *±* 0.05 (mean *±* std) seconds for 30 seconds. Beyond 30 seconds of laser excitation, the cells rounded up and were no longer motile. *Image J* [22] was used to adjust brightness and contrast of the images.

### 2.4 Image processing

#### 2.4.1 Extracting boundary fluorescence intensities

Intensity measurements of the fluorescence of myosin-GFP (myosin) and LifeAct-RFP (F-actin) were obtained using *ImageJ*. The brightness and contrast of the image was first adjusted to improve visibility. To measure intensities across all frames, a five pixel thick line segment was drawn from the cytoplasm across the actin scar and through bleb cortex (boundary of bleb) into the cell exterior. The location of the either cortex was clearly observed by a sharp increase F-actin intensity. We used the index of bleb cortex and actin scar observed along the intensity profile to extract the fluorescence intensity of both myosin and F-actin from both locations. These values were then normalized to the average intensity within the cytoplasm, henceforth referred to as the relative intensity. To minimize spatial variation in relative intensity, we averaged the measurements taken from two different locations of either cortex.

#### 2.4.2 Resetting frame rate

The frame rate for each bleb varied from 2.03 to 2.16 seconds. To obtain the average dynamics in relative intensity across all blebs, we needed measurements to be at the same frame rate. To this end, we first shifted each relative intensity time lapse to the initiation of the bleb (set to the frame before the first visible sign of a reforming cortex) and used piecewise linear interpolation. We then resampled the intensity values at the minimum frame rate of 2.03 seconds. The relative intensity measurements of 34 blebs taken from 19 cells were then averaged for further analysis.

### 2.5 Parameter estimation

All model parameters were estimated using the nonlinear least squares function within the curve fitting tool box in MATLAB R2020. To ensure that we obtained a global maximum for *n* parameters, the optimization was repeated 300 times by using initial values from an *n* dimensional Sobel set transformed to the hypercube [0, 5]^*n*^. Extending the side length of the hypercube beyond 5 did not improve the residuals of fitted parameters.

## 3 Results

We investigate the dynamics of actin and myosin in blebbing *D. discoideum* cells chemotaxing towards cyclic AMP under an agarose gel (Fig. 2a). The agarose gel helps induce blebbing by providing a pseudo-three dimensional confined environment [21]. By simultaneously detecting the cell membrane (using bright field light) and cortex (lifeact-RFP), we were able to identify the three major stages of the bleb life cycle: 1) separation of the membrane from the cortex with a visible actin scar (Fig. 2Bii, blue arrow in bright field image shows separated membrane and white arrow in lifeAct image shows absent bleb cortex. Note the absence of a detached membrane for the bright field image in Fig. 2Bi); 2) formation of a bleb cortex beneath the protruded membrane (Fig. 2Biii, with white arrow at the top of lifeAct image showing reformed bleb cortex, blue arrow at the bottom of lifeAct image showing intact actin scar and blue arrow in bright field image showing membrane) and finally 3) the loss of the actin scar with the bleb cortex still intact (Fig. 2Biv, blue arrow (bottom) showing degraded cortex, white arrow (top) showing intact bleb cortex and blue arrow on bright field showing membrane boundary). A video of the cell showing its bleb lifecycle is provided in S1 Video.

**Figure 2:**
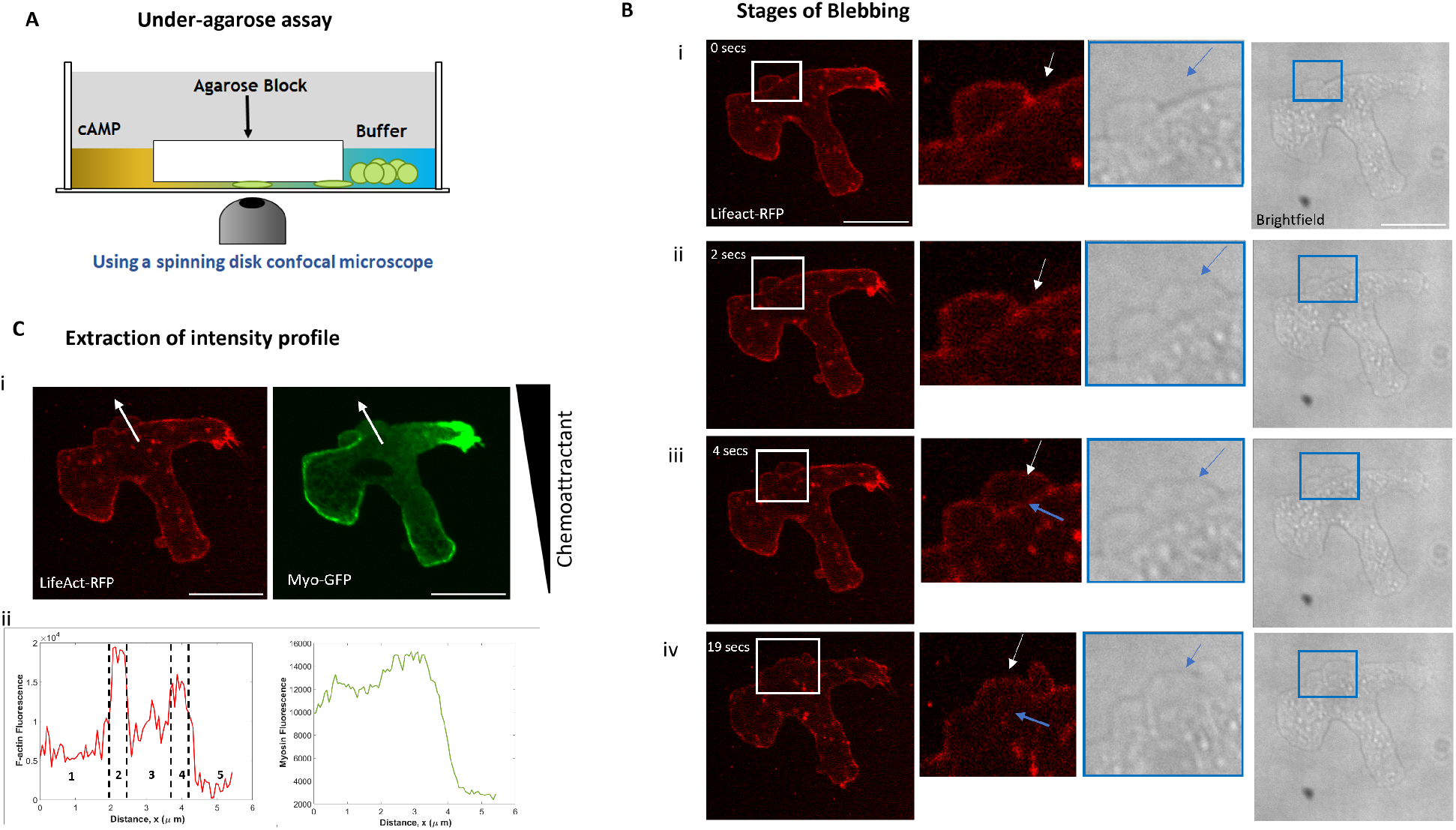
Experimental setup with stages of blebbing. A) Under-agarose assay for inducing blebs in chemotaxing *D. discoideum cells*. B) Images of F-actin (Lifeact-RFP) and the cell membrane (shown with bright field light) at different stages of blebbing. Enlarged blebbing portions of F-actin images (indicated by white box) and membrane (indicated by blue box) are shown next to the corresponding images. C) i) Representative images of blebbing cells with the line selection drawn through the bleb along which fluorescence is measured for both myosin and actin. ii) Corresponding intensity profiles shown below each image. The regions corresponding to the cytoplasm (1), actin scar (2), bleb region (3), bleb cortex (4), and cell exterior (5) are shown. Scale bar is 10 *μ*m.

*Image J* software was used to extract the fluorescent intensities of F-actin (Lifeact-RFP) and myosin (Myo-GFP) from the actin scar and bleb cortex over a duration of 30 seconds (Fig. 2C). The high intensity values in Fig. 2Cii, left correspond to the actin scar (region 2) and newly formed bleb cortex (region 4). To account for the variability in the spatial distribution of proteins along the cortex, two intensity measurements were taken from distinct locations of each bleb. The intensities are normalized to their respective cytoplasmic levels and each pair of intensities is averaged. The normalized intensities for all 34 blebs taken from 19 cells are shown for the reforming bleb cortex (Fig. 3 A1-A2) and degrading actin scar (Fig. 3 B1-B2). The results presented in this paper are based on analyzing the average relative intensities of both proteins taken over all blebs in Fig. 3(A3, B3). We note that data shown in Fig. 3 B3 was first presented in [12]. The data and figure are included in this paper for comparison to mathematical models described later in this section, which is the focus of this work. Unless otherwise specified, both F-actin and actin refer to filamentous actin for the remainder of this paper.

**Figure 3:**
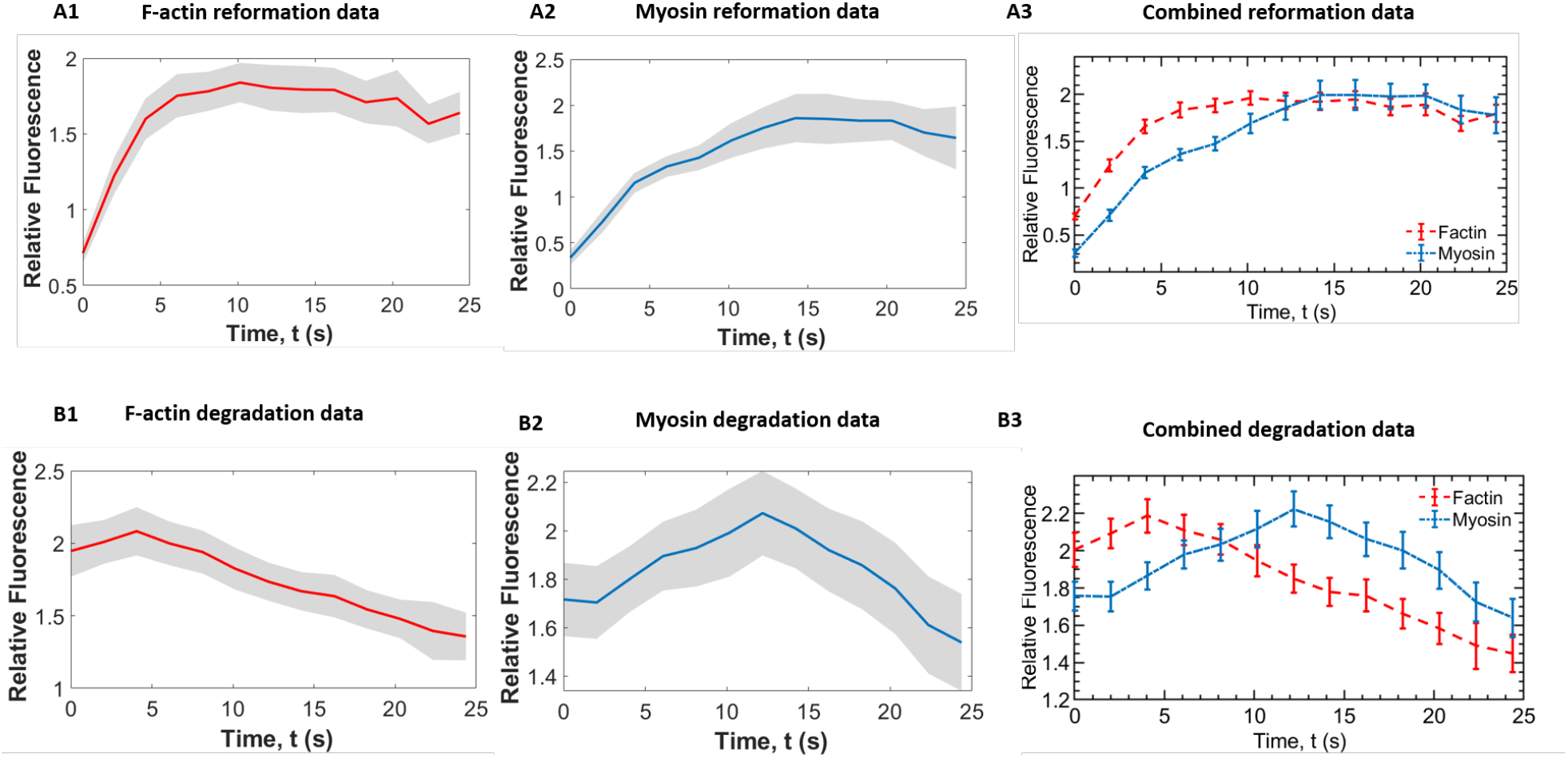
Normalized intensity profiles for complete dataset. 95% confidence interval (shaded region) and mean relative fluorescence intensities collected from 34 blebs during A1-A2) reformation of the bleb cortex and B1-B2) degradation of the actin scar. Average intensity profiles with error bars (mean *±* SEM) for A3) cortex reformation and B3) cortex degradation.

### 3.1 Experimental results of actin and myosin dynamics during reformation and degradation of the cortex

The average profiles of actin and myosin during cortex reformation reveal a rapid recruitment of actin into the bleb cortex followed by a slower recruitment of myosin (Fig. 3A3, S2 Video). Actin reaches a steady value in the bleb cortex within 5 seconds after membrane separation whereas myosin relative fluorescence levels off after about 15 seconds. Over time, the relative fluorescence of both proteins reaches a steady state value of about 1.7-1.8 times their respective cytoplasmic levels. We also observe that actin appears before myosin in a newly formed bleb (Fig. 3 A3). Fig. 4A shows that by 2.2 seconds actin can be seen within the bleb and outlining the boundary of the bleb cortex. Myosin is just emanating from the corner of the actin scar and has not yet reached the bleb cortex (white arrow and enlarged image). A similar phenomena can be observed after 6.5 seconds in a second bleb that forms after the first bleb (white arrow and enlarged image).

**Figure 4:**
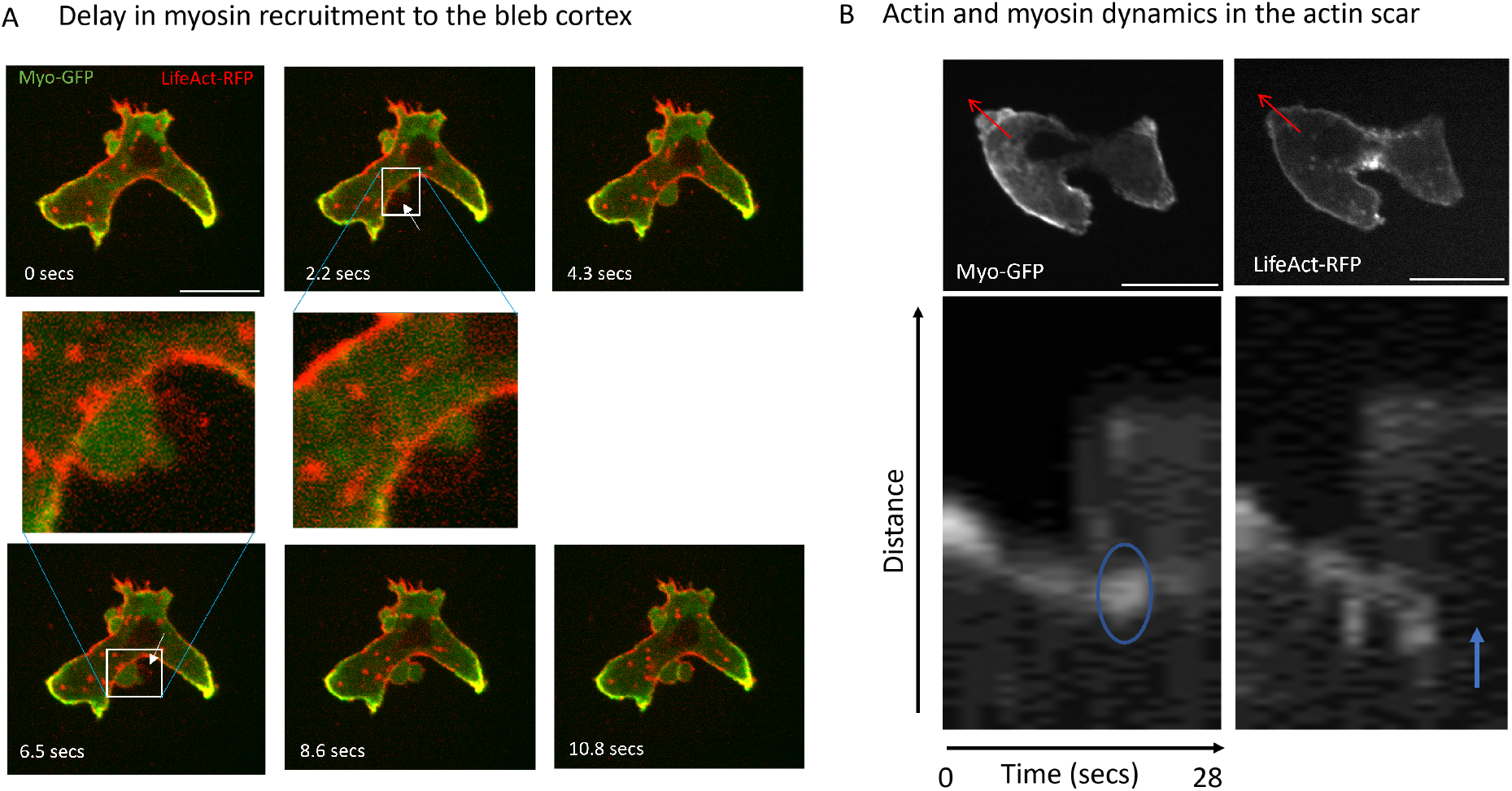
Myosin dynamics in the bleb and actin scar. A) Images of a blebbing cell with both myosin and actin channels merged to show rapid recruitment of actin to the bleb cortex and slower recruitment of myosin. Developing blebs are indicated by white arrows at time 2.2 secs and 6.5 secs with enlarged portions of the region outlined by white rectangle shown in the middle row. B) Kymographs (below) of actin and myosin intensity along a line drawn through the bleb (red arrow in cell images above). Intensity measurements are taken from the cytoplasm toward the exterior of the cell. Accumulation of myosin is seen within the blue oval in the myosin kymograph. Cortex degradation is indicated by the blue arrow in the F-actin kymograph. Scale bar is 10 *μ*m.

The degradation of the actin scar takes place over a longer time than the reformation of the cortex, lasting over 25 seconds (Fig. 3 B3). Within the first 13 seconds after membrane detachment we observe an increase in myosin intensity as the cortex is degraded, beginning from about 1.7 times its cytoplasmic intensity and reaching a maximum of 2.2 times the cytoplasmic intensity. Following this increase, the myosin intensity then declines steadily at a rate proportional to the rate at which the cortex is being degraded (Fig. 3 B3). Linear fits to the last six data points produced a slope of -0.032 (*R*^2^ = 0.9907) for actin and -0.045 for myosin (*R*^2^ = 0.9843). We generate kymographs of the myosin and actin intensity along line scans drawn through a developing bleb in Fig. 4B. Images from two additional cells are in S3 Fig. Myosin intensity increases in the region marked by the blue oval with the degraded cortex shown by the blue arrow. Fig. 3 B3 shows myosin intensity increases in the actin scar almost immediately following membrane detachment until about 12 seconds. In contrast, the degradation the actin scar occurs after about 5 seconds. Since the contractile force of myosin is known to influence actin turnover [23, 24], we hypothesize that myosin contractility may play a role in degrading actin in the scar.

### 3.2 Mathematical models for actin and myosin dynamics

We proceed to develop and analyze a family of relatively simple mathematical models for the temporal dynamics of actin and myosin during the both reformation and degradation of the cortex in blebbing cells. Our goal is to capture the essential features in the temporal dynamics of actin and myosin intensities from Fig. 3 A3 during reformation of the cortex within the bleb and from Fig. 3 B3, disassembly of the actin scar. We consider several mathematical models with different assumptions about the interaction between cortical actin and myosin, then fit the models to our experiment data.

The cortex is an active structure undergoing constant turnover due to actin polymerization and depolymerization [3]. The assembly of the cortex comprises other processes such as branching of actin filaments (F-actin) and crosslinking of branched actin filaments. Myosin binds to F-actin in the cortex when assembled into filaments (myosin filaments), which tend to form near the cortex and serve both as crosslinkers and contractile proteins [23]. Myosin filaments that unbind from actin, dissasemble and flow into the cytoplasm. We are interested in the concentration of filamentous actin *a*(*t*) and myosin filaments *m*(*t*) within the bleb cortex and actin scar, both of which are proportional to the observed fluorescent intensity from our experimental data.

For simplicity, we assume that both proteins are spatially uniform within the bleb cortex and actin scar. The concentration of F-actin in the cortex increases due to polymerization of G-actin (free actin monomeric subunits) and branching of actin filaments and decreases due to depolymerization of F-actin. Within the local blebbing region, G-actin concentration can be assumed constant since any loss of G-actin due to polymerization will be quickly replaced by diffusion of G-actin from the cytoplasm. Thus, we assume that the cortex is assembled at a constant rate 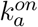. The removal of F-actin from the cortex is primarily due to depolymerization and is assumed to occur at a rate proportional to the concentration of F-actin in the cortex with rate constant 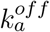. The contractile force of myosin has been reported to influence actin turnover and suggested to stabilize the actin network [24, 25]. We hypothesize that myosin influences actin turnover via the depolymerization rate, and thus investigate a myosin-dependent depolymerization rate constant 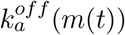. We begin from a simple assumption of a linear dependence of depolymeration on myosin concentration leading to a dissasembly rate of 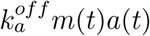. As a control for this hypothesis, we investigate a model with a myosin-independent actin depolymerization rate constant.

To model the dynamics of myosin, we assume that myosin concentration increases in the cortex at a rate that is proportional to the concentration of actin in the cortex (with binding rate constant 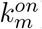) and is removed at a rate proportional to the concentration of bound myosin (with unbinding rate constant 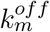). Since the cortex has a finite number of actin filaments (each with finite length), it is reasonable to assume that the binding rate constant 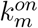 will decrease with increasing concentration of bound myosin. This leads to two competing models for the recruitment of myosin into the cortex, with a binding rate 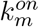 that is either dependent or independent of the concentration of myosin. Our hypotheses lead to four models for actin and myosin dynamics presented in Table 1. Model 1 assumes a myosin-independent disassembly of actin, model 2 assumes myosin-dependent actin disassembly, model 3 includes the effect of myosin on its binding to actin filaments, and model 4 includes myosin dependence for actin disassembly/depolymerication as well as myosin binding and unbinding rates.

**Table 1:** Mathematical models considered for actin and myosin dynamics during reformation of the actin cortex and disassembly of the actin scar.

| Model Name | Equations | Description | Units |
| --- | --- | --- | --- |
| Model 1 | $\frac{da}{dt} = k_a^{on} - k_a^{off} a(t)$<br>$\frac{dm}{dt} = k_m^{on} a(t) - k_m^{off} m(t)$ | Myosin-independent depolymerization | $k_a^{on} (\mu\text{Ms}^{-1}), k_a^{off} (\text{s}^{-1})$<br>$k_m^{on} (\text{s}^{-1}), k_m^{off} (\text{s}^{-1})$ |
| Model 2 | $\frac{da}{dt} = k_a^{on} - k_a^{off} m(t)a(t)$<br>$\frac{dm}{dt} = k_m^{on} a(t) - k_m^{off} m(t)$ | Myosin-dependent depolymerization | $k_a^{off} (\mu\text{M}^{-1}\text{s}^{-1})$ |
| Model 3 | $\frac{da}{dt} = k_a^{on} - k_a^{off} a(t)$<br>$\frac{dm}{dt} = \frac{k_m^{on}}{k_m + m(t)} a(t) - k_m^{off} m(t)$ | Myosin-dependent binding rate | $k_a^{off} (\text{s}^{-1}), k_m^{on} (\mu\text{Ms}^{-1})$<br>$k_m (\mu\text{M})$ |
| Model 4 | $\frac{da}{dt} = k_a^{on} - k_a^{off} m(t)a(t)$<br>$\frac{dm}{dt} = \frac{k_m^{on}}{k_m + m(t)} a(t) - k_m^{off} m(t)$ | Myosin-dependent depolymerization/binding rate | $k_a^{off} (\mu\text{M}^{-1}\text{s}^{-1})$ |

#### 3.2.1 Estimation of parameters and model selection

The variables in our models represent protein concentrations. Since our experimental data provides the fluorescence intensity of the protein in the cortex normalized to the cytoplasmic intensity, we scaled the model output by the cytoplasmic intensity of each protein, which is assumed to be constant. Denoting the cytoplasmic intensity of actin by *a*_*cyto*_ and myosin by *m*_*cyto*_ the scaled dimensionless concentrations are *a*^*^(*t*), *m*^*^(*t*). The scaled actin assembly rate and myosin unbinding rate retain their original values for all models,i.e 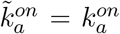 and 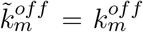. The scaled values of the remaining parameters vary with the models and are shown in Table 2. Scaled parameters in model 4 are similar to those in models 2 and 3, and therefore are not shown in the table. To simplify notation, we do not include tildes and asterisks on the model parameters and state variables, respectively, for the remainder of the paper. Results from this point forward refer to scaled variables and parameters.

**Table 2:** Scaled model parameters and their units. Scaled parameters not shown for each model have units identical to their unscaled versions.

| Model Name | Scaled Parameters | Units |
| --- | --- | --- |
| Model 1 | $\tilde{k}_m^{on} = k_m^{on} a_{cyto} / m_{cyto}$ | $s^{-1}$ |
| Model 2 | $\tilde{k}_a^{off} = k_a^{off} m_{cyto}$ | $s^{-1}$ |
| Model 3 | $\tilde{k}_m^{on} = k_m^{on} a_{cyto} / m_{cyto}^2$ | $s^{-1}$ |
| | $\tilde{k}_m = k_m / m_{cyto}$ | 1 |

We use the nonlinear least squares function within the curve fitting toolbox in Matlab to estimate the parameters of each mathematical model with objective function

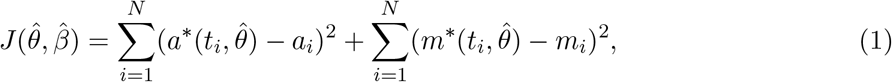

where *a*_*i*_, *m*_*i*_ denote the experimental data at time *t*_*i*_, 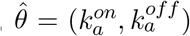 and 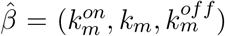 are the parameters for the scaled models. Details about the methods for sampling initial estimates of model parameters can be found in Materials and Methods. Our estimated model parameters are summarized in Table 3. Results from the nonlinear function fits and data are graphed together in Fig. 5.

**Table 3:** Values of model parameters computed from experimental data for reformation of the actin cortex in a bleb (top) and disassembly of the actin scar (bottom).

| | $k_a^{on}$<br>Assembly<br>Rate | $k_a^{off}$<br>Disassembly<br>Rate | $k_m^{on}$<br>Binding<br>Rate | $k_m^{off}$<br>Unbinding<br>Rate | $k_m$<br>Saturation<br>Constant | Residual |
| --- | --- | --- | --- | --- | --- | --- |
| Cortex reformation |  |  |  |  |  |  |
| Model 1 | 0.7602 | 0.4344 | 0.3231 | 0.3160 | NA | 0.1598 |
| Model 2 | 0.2849 | 0.0960 | 0.3007 | 0.2924 | NA | 0.1196 |
| Model 3 | 0.7603 | 0.4345 | 1.0328 | 0.2033 | 3.1566 | 0.1591 |
| Model 4 | 0.3097 | 0.1058 | 0.2433 | 0.1063 | 0.3751 | 0.0826 |
| Actin scar disassembly |  |  |  |  |  |  |
| Model 1 | $3.0882 \times 10^{-10}$ | 0.0120 | 0.1871 | 0.1705 | NA | 0.2891 |
| Model 2 | $1.1505 \times 10^{-11}$ | 0.0064 | 0.1771 | 0.1612 | NA | 0.2896 |
| Model 3 | $3.900 \times 10^{-13}$ | 0.0121 | 1.1704 | 0.1553 | 4.999 | 0.3071 |
| Model 4 | $2.8100 \times 10^{-13}$ | 0.0065 | 1.1013 | 0.1459 | 4.999 | 0.3071 |

**Figure 5:**
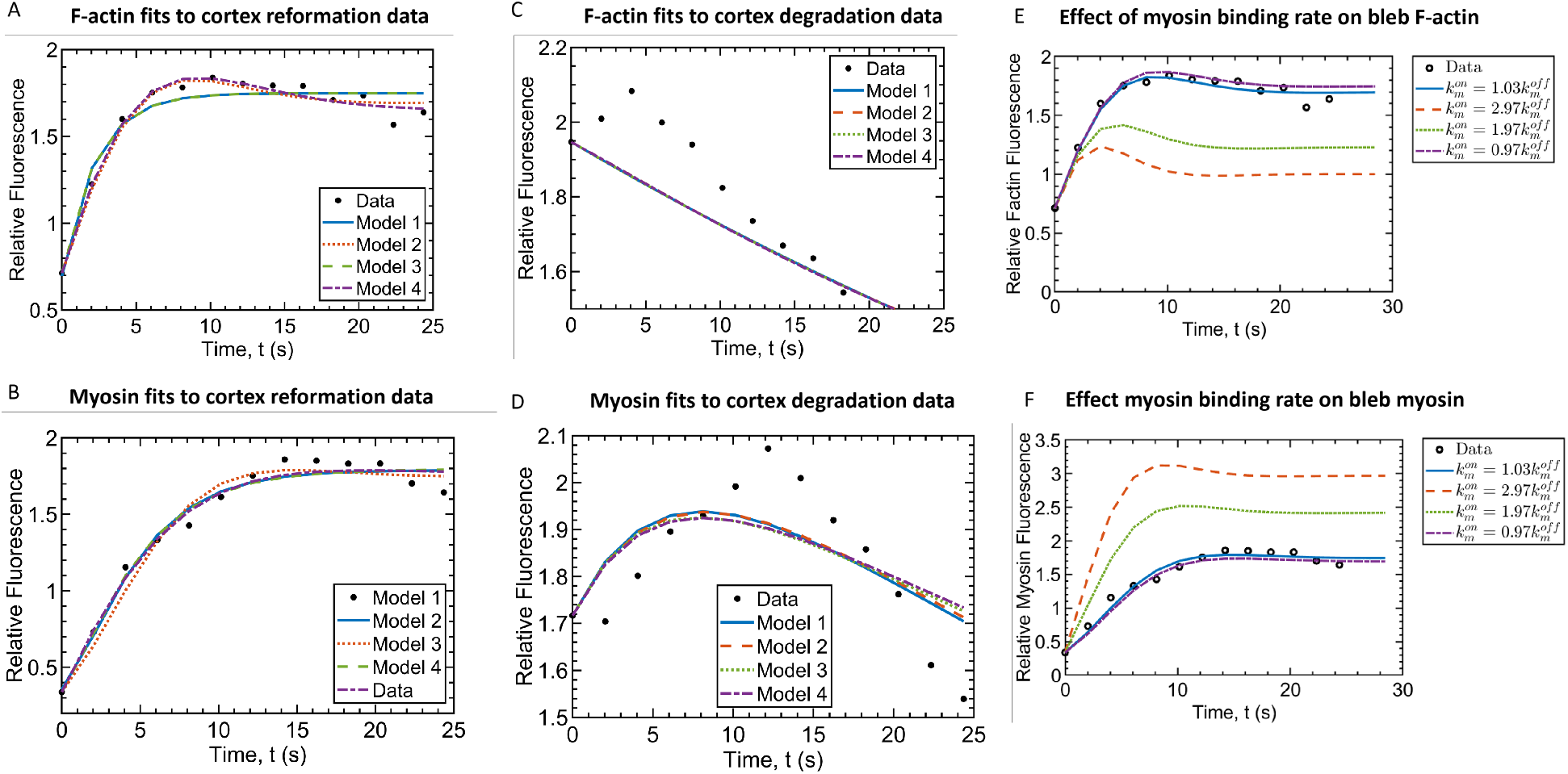
Model Fits to Experimental Data. A-B) Model fits to cortex reformation data within the bleb. C-D) Model fits to cortex disassembly data within the actin scar. (E-F) Effect of binding rate of myosin 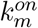 on the equilibrium concentration of actin and myosin in the cortex. The experimentally fitted accumulation rate for myosin is 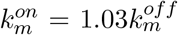 and that for F-actin is 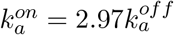.

Our estimates for actin dissasembly rate (0.1 *−* 0.43) s^−1^ are an order of magnitude larger than the turnover rates of 0.05 *s*^−1^ reported in M2 blebbing melanoma cells [26, 27]. This can be explained in part by differences in the bleb formation dynamics between *D. discoideum* and M2 melanoma cells. In particular, membrane expansion stalls after about 20 seconds in M2 melanoma cells [27] whereas the same process takes about 2 seconds in *D. discoideum* (Fig. 2B), thus requiring a faster actin turnover rate. Our estimates for myosin binding rate 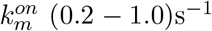 are within the range of myosin binding rates used for stochastic simulation of the actin networks at the cell periphery [28]. We note that results from the analysis to fits of the models are described in terms of the ratio of the remaining parameters.

### 3.3 Cortex reformation results

First, we focus on the fit of the models to experimental data of reformation of the actin cortex in a bleb. The fits show that models 2 and 4 achieve the lowest residual (see Table 3) and capture the dominant trends in both F-actin and myosin (Fig. 5A,B). In particular, models 2 and 4 capture the local maximum of actin during the reformation of the cortex after 5 seconds (Fig. 5a). Both of these models assume a myosin-dependent disassembly of actin filaments. The smaller residual from model 4 can be attributed in part to one additional parameter in the myosin-dependent binding rate term (first term of models 2 and 4). This conclusion is supported by the fact that inclusion of this term in model 3 did not significantly improve the fit of model 1. Thus, we consider model 2 as a sufficient model for describing the dynamics of actin and myosin during cortex reformation.

Qualitative analysis of model 2 reveals a single stable fixed point

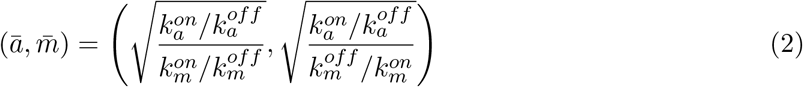

for all positive model parameters (see Appendix A for details). This suggests that cortex reformation is fairly robust to perturbations in the model kinetic parameters. From the fixed point in Eq. (2), we also observe that an increase in the effective binding rate of myosin 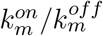, which results in a larger scaled concentration of myosin during cortex reformation will tend to decrease the equilibrium concentration of actin in the cortex, resulting in a cortex that has a lower F-actin concentration. Note that that experimental data on cortex reformation in Fig. 3A1 shows a steady state intensity of about 1.7 times its cytoplasmic value. If the effective binding rate of myosin 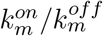 is equal to the effective accumulation rate of F-actin 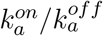, the equilibrium intensity of actin in the bleb cortex will approach its cytoplasmic concentration with relative intensity of 1, where the cortex is not well-defined (Fig. 5f). If we define a healthy cortex as one where the actin concentration is above 1.7 times its cytoplasmic concentration to distinguish cortical actin from cytoplasmic actin, then its formation requires the effective binding rate of myosin to satisfy, 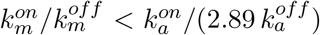. This suggests that the relatively slow accumulation of myosin is necessary to maintain a healthy, well-defined bleb cortex.

### 3.4 Cortex degradation results

The major trends in the cortex degradation data (increase in myosin followed by decrease, and a decrease in actin) are captured by all four models with very little difference between the residuals of each model (Fig. 5c-d, Table. 3). Since Model 1 assumes a myosin-independent disassembly rate and fits the experimental as well as Model 2 and 4, our modeling suggests that the effect of myosin on actin turnover is minimal when the cortex is degrading. All of the models, however, underestimate the peak in the myosin intensity data after 12 seconds and do not capture the 5 second delay in the decline in the actin intensity data (Fig. 5c-d).

Comparing the fit parameters for actin assembly and disassembly rates 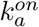 and 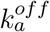 in Table 3 from reformation of the cortex to degradation of the actin scar, we observe 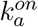 is almost zero and 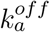 is approximately an order of magnitude smaller in the actin scar compared to cortex reformation. These results suggest that the actin assembly rate 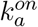 may serves as a binary switch that is turned off during the degradation of the cortex in the actin scar. When 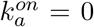, Eq. (2) shows that the system of equations from model 2 results in a zero, stable equilibrium concentration for both F-actin and myosin that is biologically relevant during disassembly/degradation of the cortex in the actin scar.

Next, we focus the models ability to fit the accumulation of myosin (local maximum) during degradation of the cortex (Fig. 3B3, 12 seconds). Since models 1 and2 both produce a similar profile for myosin during cortex degradation (Fig. 5B), and model 1 can be solved analytically, we use model 1 to identify a sufficient condition for the local maximum in myosin concentration. In order for myosin to increase initially, peak and then decrease, it is necessary for *m*(*t*) to be increasing at time zero (*dm*(0)*/dt >* 0) and have at least one positive critical point. The solutions to model 1 are

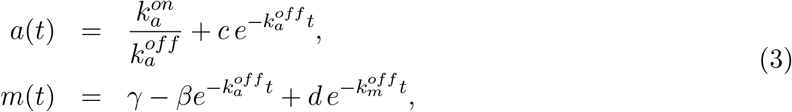

where 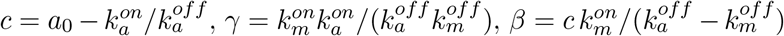 and *d* = *m*_0_ + *β − γ*. The myosin concentration increases initially as long as 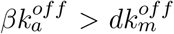, which can be simplified to the condition

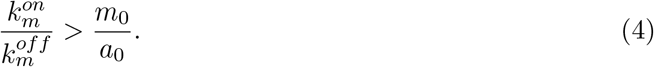

The myosin solution *m*(*t*) a single critical point at

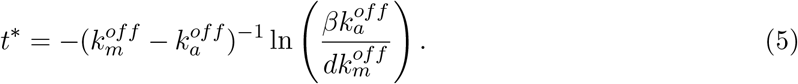

*m*(*t*^*^) is positive if

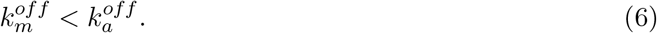

Details on the solution process and analysis of the conditions for the local maximum in *m*(*t*) are provided in Appendix B. In summary, our analysis suggests that a local maximum in myosin in a degrading cortex occurs if the myosin unbinding rate is smaller than the disassembly rate of the cortex (Eq. (6)) and the ratio of myosin binding to unbinding rate is sufficiently large (Eq. (4)). Indeed, our estimated parameters satisfy these conditions (see Table 3).

### 3.5 Varying the actin assembly rate

Our model fit to the cortex degradation intensity data does not capture the local maximum in the F-actin solution at 5 seconds (Fig. 5C). The estimated near zero polymerization rate captures the average rate of decline of the cortex after this delay period, suggesting that this parameter is more relevant at the onset of degradation and not immediately following membrane separation. The transition between a non-degrading cortex and an actively degrading cortex with decreasing F-actin intensity/concentration suggests a the actin assembly rate 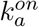 may not be constant. We tested this idea by adjusting model 2 to include a assembly rate that decreases exponentially with time, i.e. 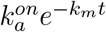. This formulation significantly improved our model fit to both the cortex reformation 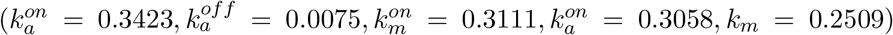 and degradation data 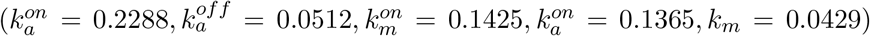 with residuals of 0.0695 and 0.0674, respectively. Experimental intensity data and model function fits are shown in Fig. 6.

**Figure 6:**
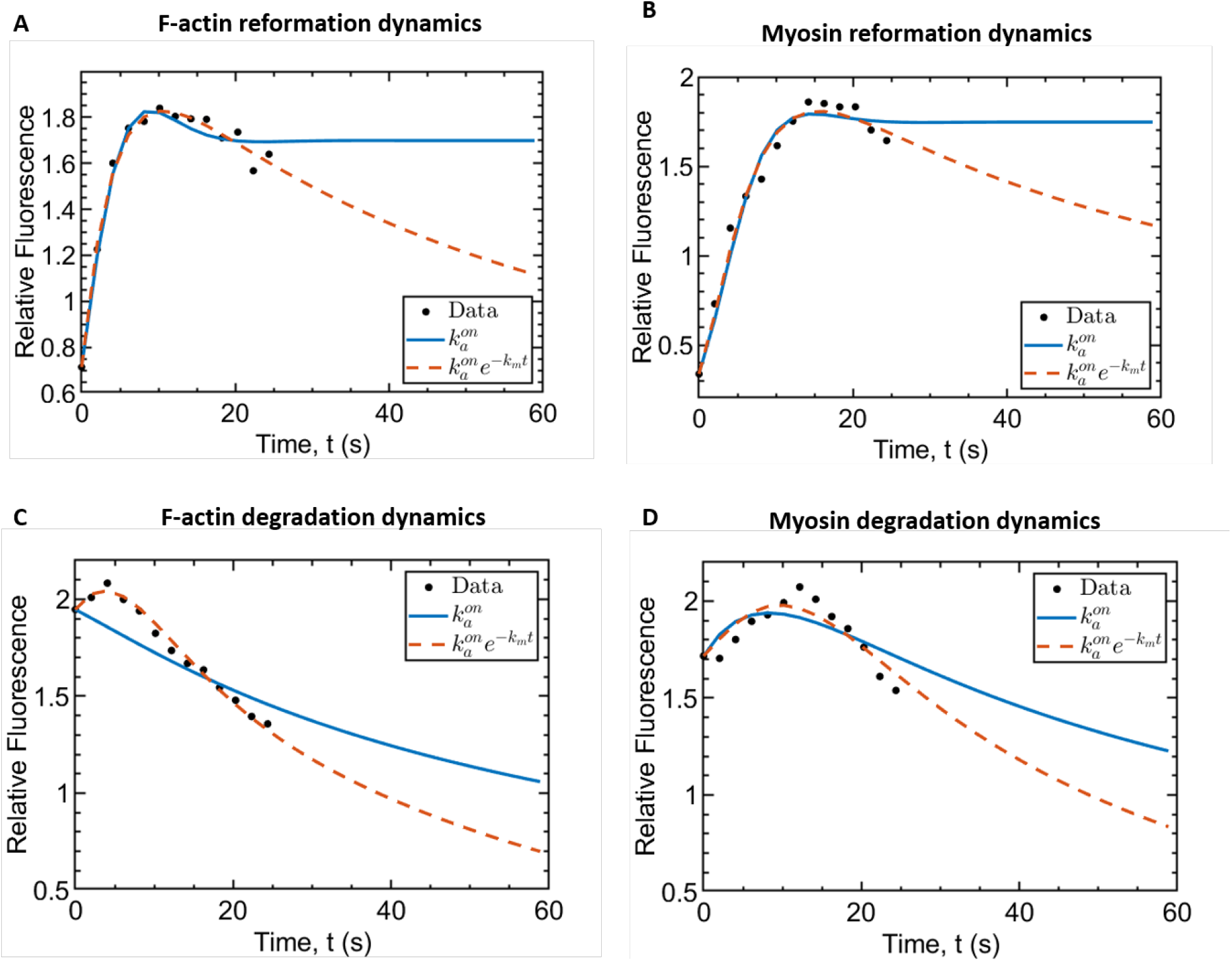
Fit and long term dynamics of alternative model for actin polymerization. Model fits and long term dynamics for A-B) cortex reformation data and C-D) cortex degradation data using the exponentially decreasing polymerization rate.

While the model with exponentially decreasing polymerization rate fit the cortex reformation data well, it did not capture the long term behavior of the reforming cortex. For both reformation and degradation of the cortex, the actin intensity decreased to zero (Fig. 6A,C). However, the long term behavior for cortex degradation data approaches zero (Fig. 6C,D), which is expected biologically. We expect the reforming cortex to approach a value of 1.7 to maintain a well-defined cortex.

## 4 Discussion

### 4.1 Dynamics of actin and myosin during bleb stabilization differs from retraction

In this work, we examine the dynamics of actin and myosin in the actin scar and bleb cortex during bleb-driven motility in chemotaxing *D. discoideum* cells. Our analysis of fluorescence intensity data from F-actin and myosin reveal a cortex reformation timescale of about 5 seconds that is much faster than the timescale of cortex degradation (over 25 seconds). Our data shows that the actin scar does not degrade during the first 5 seconds of active cortex reformation. During cortex reformation myosin intensity lagged behind F-actin intensity by about 2 seconds. Our results on the rapid accumulation rate of F-actin in the developing bleb cortex and the slower pace of myosin accumulation are consistent with actomyosin assembly dynamics observed in blebs isolated from HeLa and M2 cells (see [2], Fig. 4c-d). Whereas blebs retracted in nonmigrating HeLa and M2 cells studied in [2] and F-actin and myosin concentration continued to increase after the bleb achieved its maximal size, we observe that in migrating cells both proteins reach a steady-state within 15 seconds. These observations reveal a critical difference in dynamics of myosin and F-actin when blebs are stabilized for movement compared to when they are retracted back into the cell body.

### 4.2 Contractile force of myosin supports the stabilization of the bleb cortex

Simulation of our mathematical model (model 2) reveals that a high accumulation rate of myosin in the bleb cortex tends to decrease the equilibrium concentration of actin (Fig. 5d). In particular, we show that if myosin accumulates at the same rate as actin, no cortex will be formed. Our observations and theoretical simulations are supported by recent experiments on retracting blebs where myosin accumulation was associated with a relaxation in the cortex density and a return of the bleb cortex density to the actin scar density [2]. The delay in myosin’s recruitment is also supported by a related study in filamin-deficient M2 cells where contractile proteins were among the last proteins to be recruited to the reforming cortex [9]. Our mathematical model (model 2) suggests that myosin’s influence on the concentration of actin in the bleb cortex is through regulation of the disassembly rate of actin, as evidenced by a relatively small residual for the fit (see Table 3). This mechanism of myosin regulated disassembly is supported by recent observations that sub-piconewton pulling forces applied in vitro to Arp 2/3-mediated filament branches increases their detachment from the mother filament [29, 30]. Also, a related study involving gliding assays with branched filaments on motor protein surfaces has shown that pulling forces from motors could cause actin filament debranching [31, 29].

### 4.3 Membrane separation triggers the degradation of the actin scar by decreasing the rate of actin assembly

Our model suggests that the degradation of the actin scar can be explained by an exponential decrease in the assembly rate of actin. This is in contrast to the constant assembly rate that resulted in good model fits to intensity data from the reformation of the cortex. A major physical differences between the two processes (reformation and degradation) is that cortex reformation occurs within the proximity of the membrane whereas cortex degradation occurs in the region devoid of the cell membrane. This suggests a role for membrane position on the reformation or degradation of the cortex.

The assembly of the cortex which comprises actin polymerization and branching of actin filaments is regulated by several proteins, with Arp 2/3 being one of the major proteins regulating branching. Arp 2/3 is activated by nucleation-promoting factors (NPFs) which includes the WASP and WAVE family of proteins [32]. These proteins are anchored to the lipid membrane, where they help to bring together Arp 2/3 and actin filaments to facilitate the formation of branched actin [32, 29, 33]. NPFs also help to speed up the growth of actin filaments by delivering actin and profilin-actin to the barbed ends of growing filaments [29]. In vitro experiments have shown that the branched actin networks growing against an artificial NPF-decorated surface can elongate faster than barbed ends in solution [34]. Thus, it stands to reason that the separation of the membrane the from the actin scar, which removes NFPs from the bleb cortex, will decrease its assembly rate. First, through a cessation of branched actin formation and then by slowing down the elongation of filaments. Our theoretical study suggests that this decrease in the actin assembly rate is not instantaneous but rather occurs exponentially (see Fig. 6c). Conversely, the availability of the cell membrane at the boundary of the bleb and G-actin within the cytosol facilitates the formation of a new branched actin network. Model 2 suggests that this reformation process occurs at a constant assembly rate (see Fig. 6a).

### 4.4 Membrane separation triggers the accumulation of myosin into the degrading cortex

Our experimental data show a correlation between myosin accumulation and the degradation of the actin scar. In particular, we observe an accumulation of myosin while the cortex degrades in the actin scar. However, our theoretical analysis did not identify a significant role for myosin in this process. In particular, myosin concentration did not significantly influence the disassembly rate of the cortex. This is surprising in light of several studies that suggested myosin may increases actin filament debranching by pulling on them [29, 30, 35]. Other studies have also pointed to myosin contractility as a driver of cortex breakage which helps to initiate blebbing [8, 36]. While it is still possible for the accumulation of myosin to help dissasemble the degrading cortex, our study finds that the modulation of actin assembly rate plays a more significant role.

In spite of the seeming minor role played by myosin accumulation (local maximum) in cortex degradation, we sought to understand potential regulators of this process. Analysis of model 1 revealed that a high effective binding rate of myosin (Eq. 4) and an unbinding rate which is smaller than the actin disassembly rate (Eq. 6) are necessary for myosin to accumulate in a degrading cortex. Any factor that decreases the unbinding rate of myosin 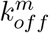 would simultaneously meet both accumulation conditions. The unbinding rate of myosin is promoted by Myosin Heavy Chain Kinase A (MHCKA)[37], whose activation is regulated by a cyclic AMP induced signaling pathway which begins with the transmembrane receptor cAR1 [38]. Thus, separation of the membrane from the cortex could drive the observed accumulation of myosin into the degrading cortex.

### 4.5 Model limitations

Here, we consider two species systems of differential equations for actin and myosin with relatively simple kinetic rate laws. There are many proteins involved in forming the F-actin cortex, including ARP 2/3, formins, crosslinking proteins, capping proteins, turnover proteins, etc. [5, 2]. Our motivation for considering such simple models comes from our experimental data being limited to only actin and myosin relative fluorescence data. In spite of the simplicity of the models, they result in good fits of the cortical reformation data over time (Fig. 5A,B). The model fits to actin and myosin relative fluorescence data during degradation of the actin scar are not as good, unless we include an exponentially decreasing actin assembly rate (compare Fig. 5C,D to Fig. 6C,D). Since our results did not identify a significant role for myosin in degradation of the actin scar, other factors such as actin-depolymerizing factor (ADF)/cofilin could be important in degrading the actin scar. A more detailed model would include such terms as well as well as experimental data for additional proteins.

## Author Contributions

**E.O Asante-Asamani**: Conceptualization, Formal analysis, Investigation, Methodology, Visualization, Writing-original draft, Writing-review & editing. **Mackenzie Dalton:** Visualization, Formal analysis. **Derrick Brazill:** Funding acquisition, Investigation, Writing-review & editing. **Wanda Strychalski:** Conceptualization, Formal analysis, Investigation, Methodology, Visualization, Writing-review & editing.

## Acknowledgements

This work was supported by grants to D-B. from the National Science Foundation (MCB-1244162), a PSC-CUNY grant (692710047), as well as Research Centers in Minority Institutions Program grants from the National Institute on Minority Health and Health Disparities (8 G12 MD007599) from the National Institutes of Health. E-A.A is supported by the AMS Simons Travel Grant. This work was also supported in part by grant #429808 from the Simons Foundation to WS.

### Appendices

#### A Fixed point and local stability of mathematical model

Our mathematical model for actin and myosin dynamics during cortex reformation (model 2), as presented in Table. 2 in the main manuscript is

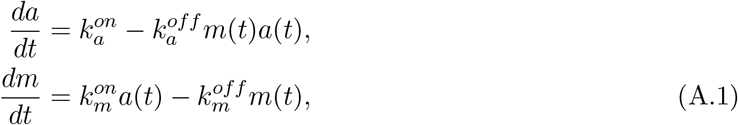

where we do not include the asterisk on actin and myosin concentration to simplify the notation. The fixed point of the model (*ā*, 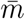) satisfy the equations 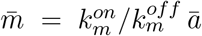 and 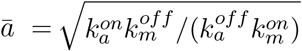, yielding the value

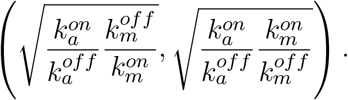

Linearizing the nonlinear model (A.1) we obtain the Jacobian matrix

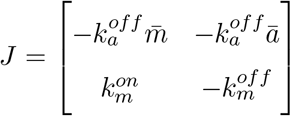

with trace 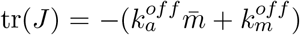 and determinant, 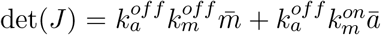. Since the trace is always negative, the fixed point is always stable.

#### B Sufficient condition for myosin accumulation

Here, we offer details to support the sufficient condition for which myosin can be expected to accumulate, or have a local maximum, in a degrading cortex. Our analysis is based on the linear model for actin and myosin dynamics (model 1) which is

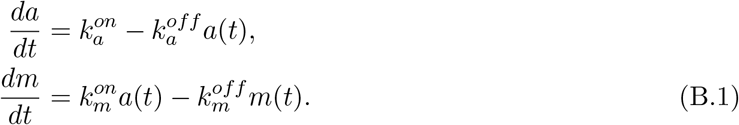

Since actin concentration equation is decoupled from the myosin equation, we solve this exactly using the method of integrating factors to obtain the exact solution

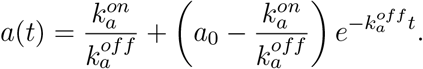

Substituting this into the myosin equation, we solve the resulting differential equation using an integrating factor,

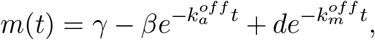

where 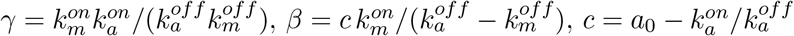, and *d* = *m*_0_ + *β − γ*. In order for myosin to initally increase, achieve a local maximum, and then decrease, it is sufficient for

the solution to initially increase, *m*^l^(0) *>* 0 and for there to be a single positive critical point. From the derivative of the myosin solution, 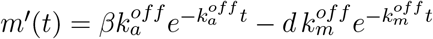, we find that *m*^l^(0) *>* 0 whenever

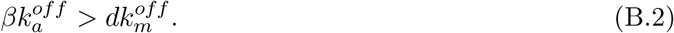

The solution also has a single critical point at *t*^*^ satisfying

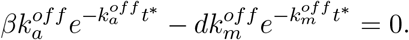

Solving the above equation yields 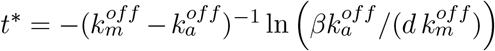. Since we require that 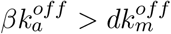 for myosin to initially increase, the additional condition for the critical point to be positive is 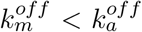. Therefore, if a) 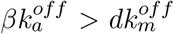 and b) 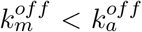, myosin in the cortex will accumulate at a positive time value *t*^*^. Recall that 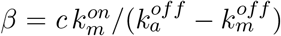, *d* = *m*_0_ + *β − γ*, 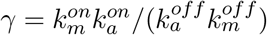 and 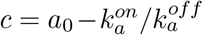. Condition a) together with the definition of *d* implies that

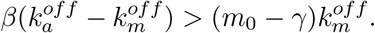

Substituting the definition for *β* and *γ* and simplifying the resulting inequality we have that

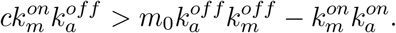

Now substituting the definition for *c* and simplifying, condition a) reduces to the requirement that

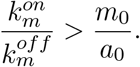

Thus the two conditions for myosin to accumulate in a degrading cortex are:

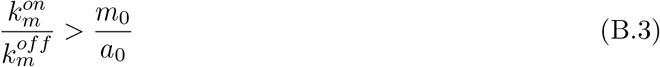

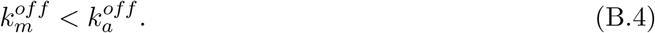

## Supporting Information Captions

**S1 Video. Bleb formation cycle**. The video shows intensity of F-actin (LifeAct RFP) and membrane (Brighfield) during the bleb formation cycle for the cell in Fig. 2

**S2 Video. Delay in myosin recruitment during bleb formation** Video of cell in in Fig. 4 showing a delay in myosin recruitment to the bleb cortex.

**S3 Fig. Evidence of myosin accumulation**. Kymographs of two additional cells showing an accumulation of myosin in the actin scar during cortex degradation.

