## Supplementary figures and images for "Modeling the dynamics of actin and myosin during bleb stabilization"

### Supplemental Figure 3

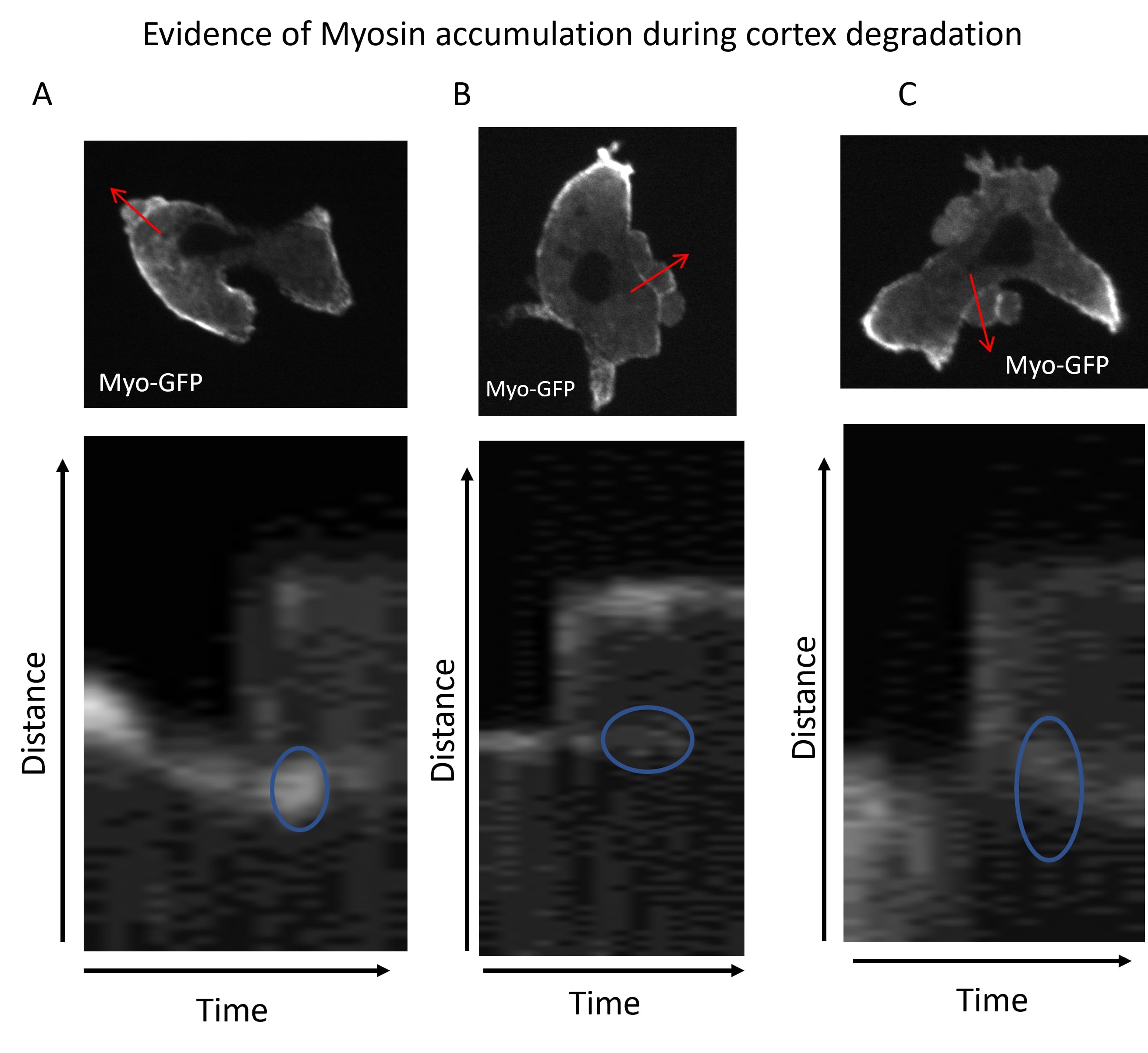
